# A comprehensive landscape of 60S ribosome biogenesis factors

**DOI:** 10.1101/2021.05.11.443624

**Authors:** Carolin Sailer, Jasmin Jansen, Jan P. Erzberger, Florian Stengel

**Affiliations:** Department of Biology, University of Konstanz, Universitätsstraße 10, 78457 Konstanz, Germany; Konstanz Research School Chemical Biology, University of Konstanz, Germany; Department of Biophysics, UT Southwestern Medical Center - ND10.214C, 5323 Harry Hines Blvd., Dallas, TX 75390-8816, USA

**Keywords:** ribosome biogenesis, ribosome biogenesis factors (RBFs), ribosome assembly factors (AFs), crosslinking coupled to mass spectrometry (XL-MS), mono-intralink filter (mi-filter), false discovery rate (FDR), DEAD-box helicases

## Abstract

Eukaryotic ribosome biogenesis is facilitated and regulated by numerous ribosome biogenesis factors (RBFs). High-resolution cryo-EM maps have defined the molecular interactions of RBFs during maturation, but many transient and dynamic interactions, particularly during early assembly, remain uncharacterized. Using quantitative proteomics and crosslinking coupled to mass spectrometry (XL-MS) data from a extensive set of pre-ribosomal particles, we derived a comprehensive and time-resolved interaction map of RBF engagement during 60S maturation. A novel filter that efficiently eliminates false positive interactions and integration of our MS data with known structural information allowed us to localize 22 unmapped RBFs to specific biogenesis intermediates and to identify 9 proteins that represent potentially new RBFs. Our analysis reveals an extensive interaction network for the casein kinase complex in 60S maturation and elucidates the timing and molecular function of 60S engagement by DEAD-box ATPases. Our data provide a powerful resource for future studies of 60S ribosome biogenesis.

## Introduction

Ribosome biogenesis is a complex, multistep maturation process that assembles protein and RNA components across multiple cellular compartments to generate mature ribosomes. Around 200 ribosomal biogenesis factors (RBFs) engage pre-ribosomes during assembly but are not components of mature ribosomes (Klinge and Woolford, 2019; Konikkat and Woolford, 2017; Kressler et al., 2017; Woolford and Baserga, 2013). The majority of ribosomal RNA (rRNA) is synthesized as a single transcript, the 35S rRNA, that contains the 18S (small subunit) as well as the 5.8S and 25S rRNA (large subunit) elements found in mature ribosomes. RBFs are crucial for the precise processing of 35S rRNA into mature rRNAs and for the ordered incorporation of ribosomal proteins (r-proteins) during biogenesis. RBFs achieve this by engaging specific intermediate states and by guiding structural and conformational rearrangements of the rRNA during maturation. Changes in RBF composition correlate with the movement of pre-ribosomes from the nucleolus to the cytoplasm, suggesting that RBFs composition guides the cellular movement of pre-ribosomes. Depending on their individual functions, some RBFs engage maturing ribosomes transiently, while others remain bound for prolonged periods during biogenesis. The focus of this study is the 60S biogenesis pathway in the yeast *Saccharomyces cerevisiae,* beginning with intermediates that immediately precede the physical separation of 18S and 27S rRNA after 35S cleavage at the A2 site within the internal transcribed spacer 1 (ITS1). Initially, studies of 60S biogenesis focused on rRNA processing events, especially the precise, multi-step excision of internal transcribed spacer 2 (ITS2), located between the 5.8S and the 25S rRNA on the 27S pre-rRNA, and on the proteomic characterizations of various affinity purified biogenesis intermediates (Gamalinda et al., 2014; Woolford and Baserga, 2013). More recently, multiple cryo-EM reconstructions of 60S pre-ribosomal particles have allowed the structural mapping of many assembly factors and offered insights into their hierarchical incorporation in sequential 60S intermediates (Kargas et al., 2019; Kater et al., 2020; Kater et al., 2017; Sanghai et al., 2018; Wu et al., 2016; Zhou et al., 2019a; Zhou et al., 2019b).

Despite this remarkable assortment of structural models, the low abundance and inherent dynamics of pre60S assemblies still make it extremely challenging to resolve structures of more transient assembly intermediates or to visualize dynamic regions, something that is particularly true for early nucleolar assembly intermediates. This is reflected in the fact that no structural or functional information exists for over half of the known RBFs associated with 60S biogenesis, even though they play important roles in 60S maturation.

Among ribosome biogenesis factors whose structural role has remained elusive are seven members of the DEAD-box family of ATPases. These proteins are thought to play fundamental roles in ribosome biogenesis, catalysing energy-consuming RNA remodelling events to unidirectionally drive specific maturation steps during ribosome assembly (Klinge and Woolford, 2019; Martin et al., 2013; Rodríguez-Galán et al., 2013). To date, of the eight DEAD-box helicases associated with 60S biogenesis, only Has1 has been successfully positioned in high-resolution cryo-EM maps of 60S pre-ribosomal particles (Kater et al., 2017; Sanghai et al., 2018; Zhou et al., 2019a).

In this study, we use large scale biochemical enrichments of a comprehensive set of 60S pre-ribosomal particles in the yeast *Saccharomyces cerevisiae* to identify and quantify individual RBF abundances within 60S intermediates and use them to create a comprehensive and precise timeline of RBF engagement during 60S maturation. In addition, XL-MS was used to identify interaction partners for individual RBFs, implementing a novel mono- and intralink filter (mi-filter) that significantly improves false-discovery rates for medium to large scale XL-MS data. Combined, this data allows us to expand the 60S engagement profile of structurally characterized factors, to localize structurally uncharacterized RBFs and to identify potential novel 60S biogenesis factors. Overall, our comprehensive data represents an essential resource for future structural studies of large subunit ribosome biogenesis

## Results

### An overall timeline of 60S biogenesis derived from quantitative MS-data

To generate a comprehensive, time-resolved dataset of RBF engagement during 60S biogenesis, we selected RBFs covering the entire 60S maturation pathway as baits for the large-scale biochemical enrichment of 60S pre-ribosomal particles. Samples were analysed using our integrated Affinity-Purification Label-Free Quantification and XL-MS (AP-MS LFQ/ XL-MS) workflow (**Figure 1A**). This strategy allowed us to simultaneously generate a precise proteomic profile of RBF abundance by identifying and quantifying non-crosslinked peptides in each sample, thereby reconstructing the overall timeline of 60S ribosomal biogenesis, and to define the protein interaction network for each of the purified pre-ribosomal particles using chemical crosslinking (**Figure 1A**). In total, the affinity purification of 36 independently purified samples from 12 different RBFs allowed us to reliably quantify 272 proteins using MaxQuant (Hein et al., 2015; Tyanova et al., 2016a) in combination with the Perseus package for statistical validation (Tyanova et al., 2016b) (**Figure 1B to D**, **Figures S1 to S4**, **Supplementary Data 1**; *see* STAR Methods for details).

**Figure 1:**
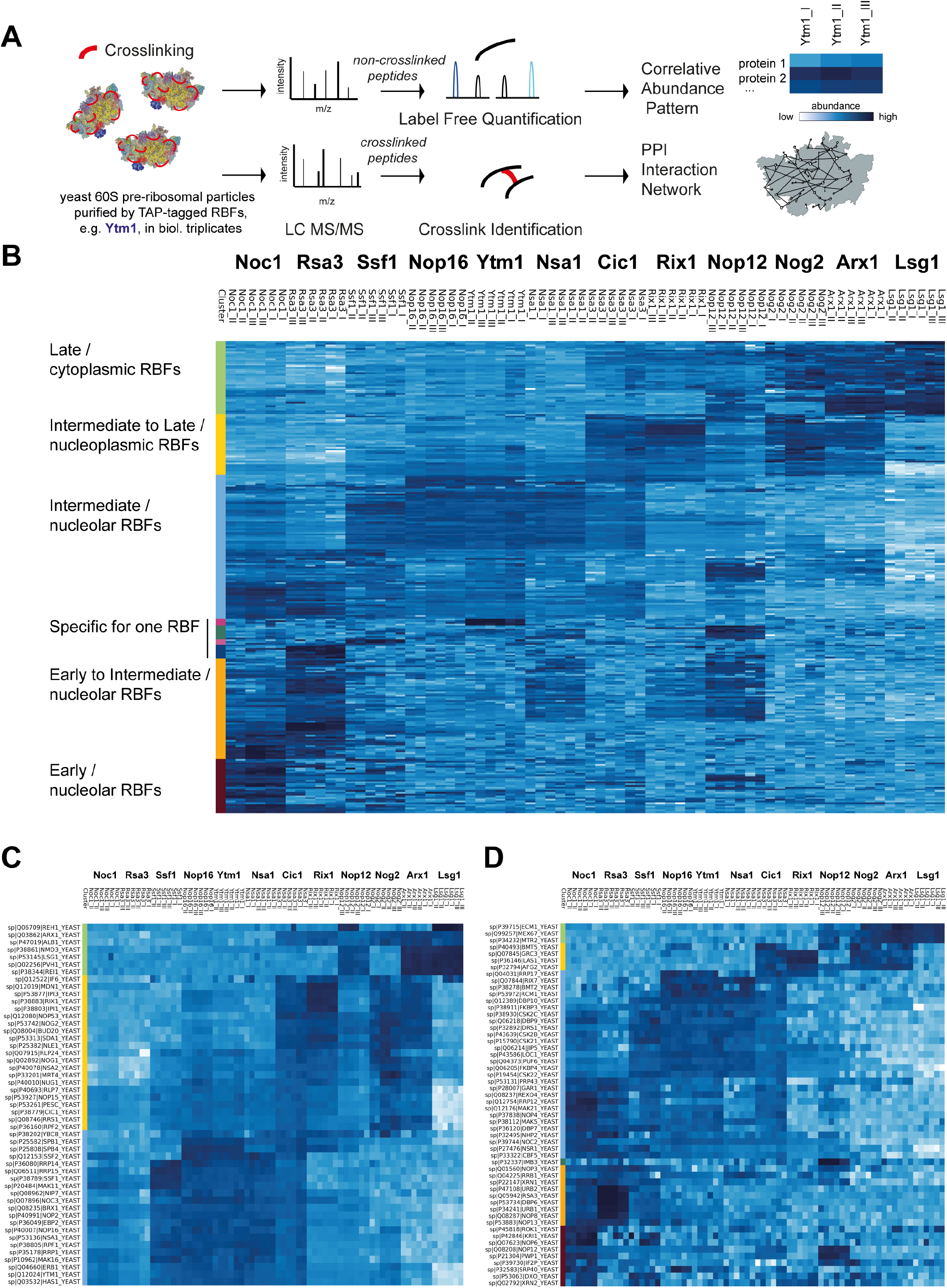
Timeline of 60S ribosomal biogenesis. (**A**) Schematic overview of our AP-MS LFQ & XL-MS workflow. Different RBFs were used to affinity purify 60S pre-ribosomal particles. Particles were crosslinked and analysed by LC-MS/MS. Non-crosslinked peptides were used for the identification and quantification of proteins with a label free approach in order to generate a correlative abundance pattern and crosslinked peptides were used to obtain particle specific protein-protein interaction networks. (**B**) Abundance pattern of quantified proteins from AP-MS data used to generate a timeline of RBF involvement in 60S ribosome assembly. Shown is a heatmap of all 272 proteins, which were reliably identified and quantified in pre-ribosomal particles from 12 different RBFs used as bait proteins (biological triplicates; n=3). RBFs used as bait proteins (x-axis) are shown on top and are plotted versus their respective interactors (y-axis). Protein abundances are sorted by unbiased hierarchical clustering and are shown as Z-score normalized log2 transformed LFQ values ranging from ≤ −4.0 (white) to 0.0 (azure blue) to ≥ 2.5 (dark blue) from early to late assembly states. Interactors are additionally grouped into ‘early’ (red), ‘early-intermediate’ (orange), ‘intermediate’ (blue), intermediate-late’ (yellow) and ‘late’ (green) proteins. (**C**) Abundance pattern of RBFs with known localization from previous pre-60S high-resolution studies. (**D**) Abundance pattern of RBFs for which no structural information at pre-ribosomal particles has been available so far.

Unbiased hierarchical clustering based on similarities in measured protein abundances shows that biological replicates always cluster together, reflecting the robustness and reproducibility of our sample purification and analysis workflow (**Figure S1**). The clustering analysis places 60S intermediates purified with affinity-tagged Noc1 at the beginning of the timeline, followed by intermediates containing Rsa3, Ssf1, Nop16, Ytm1, Nsa1, Cic1, Rix1, Nop12, Nog2, Arx1 and Lsg1 (**Figure S1**). This timeline is broadly consistent with previous structural, biochemical and genetic studies (Klinge and Woolford, 2019; Kressler et al., 2010, 2017; Woolford and Baserga, 2013). Nine of our baits (Ssf1, Nop16, Ytm1, Nsa1, Cic1, Rix1, Nog2, Arx1 and Lsg1) have been identified and modelled in cryo-electron microscopy (Cryo-EM) reconstructions of 60S intermediates (Kargas et al., 2019; Kater et al., 2020; Kater et al., 2017; Sanghai et al., 2018; Wu et al., 2016; Zhou et al., 2019a; Zhou et al., 2019b) and their distribution among known 60S intermediate structures is consistent with the timeline derived from our MS data (**Figure 2A**). Our other bait RBFs (Noc1, Rsa3 and Nop12) have not been identified in any 60S structural reconstructions to date. In line with previous proteomic studies (de la Cruz et al., 2004; Milkereit et al., 2001), our analysis places Noc1 and Rsa3 at the early end of our timeline, with a protein distribution pattern that is distinct from later intermediates (**Figure 1B**). In particular the presence of the RBFs Prp5, Rok1 and Rex4, critical mediators of ITS1 processing (Eppens et al., 2002; Khoshnevis et al., 2016), other RBFs associated with small subunit biogenesis as well as 40S r-proteins, suggest that these intermediates span the cleavage of 35S into small and large ribosomal rRNA fragments, and thus represent the earliest independent pre-60 complexes. Intermediates purified using affinity-tagged Nop12 cluster with nucleoplasmic intermediates containing Nog2 and Rix1 in our overall timeline of 60S ribosome biogenesis. However, the pattern of particles purified with affinity-tagged Nop12 shows additional complexity and some enrichment with proteins linked to early or early to intermediate nucleolar 60S assembly stages (**Figure 1B to D, S2 and S3**), thus not ruling out a broader involvement of Nop12 in 60S biogenesis.

**Figure 2:**
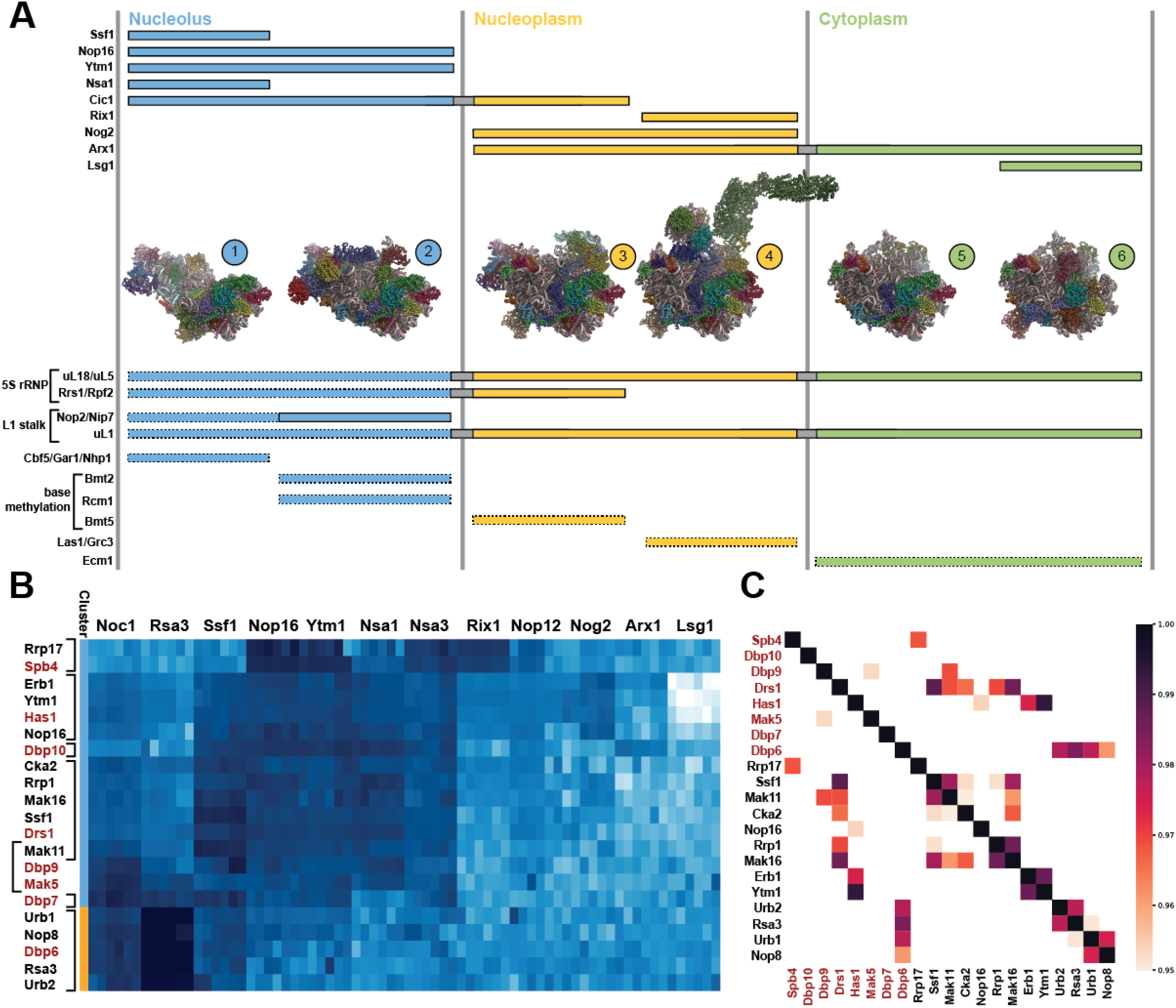
Assigning structurally uncharacterized RBFs to specific 60S cryo-EM reconstructions. (**A**) TOP – distribution of affinity-tagged RBFs used for our timeline among representative models derived from pre-60S cryo-EM reconstructions. MIDDLE-nucleolar pre-60 intermediates:1 (pdb 6EM1), 2 (pdb 6ELZ); nucleoplasmic pre-60 intermediates: 3 (pdb 3JCT), 4 (pdb 6YLH); cytoplasmic pre-60 intermediates: 5 (pdb 6N8J), 6 (pdb 6RZZ). BOTTOM –extended distribution of 5S rRNP (uL5, uL18, Rpf2 and Rrs1) and L1 stalk (Nop2, Nip7 and uL1) complexes to early nucleolar intermediates based on quantitative mass spec data (dashed boxes) and association of rRNA modifying enzymes to distinct pre-60S structural intermediates derived from correlation with quantitative mass spectrometry data. Colour patterns are the same as in Figure 1. (**B**) Heatmap of abundance patterns of DEAD-box ATPases in our timeline, along with most closely associated factors as determined by Pearson correlations. Protein abundances are shown as Z-score normalized log2 transformed LFQ values ranging from ≤ −3.0 (white) to −0.5 (azure blue) to ≥ 2.0 (dark blue). Close associations of DEAD-box proteins (highlighted in red) are bracketed. (**C**) Heatmap of pairwise Pearson correlations of the proteins shown in panel B. DEAD-box proteins are labeled in red. Correlations >0.95 are shown and coloured according to the legend.

Our final dataset contains a total of 131 proteins previously annotated as ribosome biogenesis factors in the literature (Woolford and Baserga, 2013) or in the *Saccharomyces* genome database (SGD) (Cherry et al., 2012). In total, 81 RBFs have been modelled at atomic resolution into cryo-EM maps of pre-40S (28 RBFs) and pre-60S (53 RBFs) ribosomal intermediates, of which we find 77 in our dataset (Barandun et al., 2017; Cheng et al., 2017; Kargas et al., 2019; Kater et al., 2020; Kater et al., 2017; Kornprobst et al., 2016; Schuller et al., 2018; Wu et al., 2016; Zhou et al., 2019a). The presence of pre-40S RBFs in the early stages of our timeline indicates that some of our earliest pre-60S samples are still tethered to pre-40S particles via ITS1 and that our samples therefore encompass the entire 60S assembly landscape. We reliably identify and quantify 49 of the 53 RBFs modeled into cryo-EM maps of 60S pre-ribosomal particles, (the missing factors are Cgr1, Ria1, Sdo1 and Rtc3) (**Figure 1C**, **Figure S3, Supplementary Data 1**). The near complete coverage of previously localized RBFs and the accurate clustering of characterized RBFs validate our approach and speak to the high reliability of our dataset. In addition to the 49 RBFs with established positions in pre-60S particles, we identify and quantify an additional 47 (of the remaining 54) proteins in our dataset that have been linked to large subunit biogenesis through biochemical and genetic studies (Cherry et al., 2012; Woolford and Baserga, 2013) (**Figure 1D**, **Supplementary Data 1**).

Peptides of an additional 82 non-ribosomal proteins can be readily detected and quantified in our samples (**Figure S4, Supplementary Data 1**). The known functions, high cellular abundance and lack of an obvious link to ribosome biogenesis suggest that most of these proteins are minor contaminants in our purifications. However, a subset of factors are potentially uncharacterized biogenesis factors, including Tma16, Stm1, Nap1, Lhp1, Pab1, Sro9, YMR310C, YGR283C and YCR016W. Tma16 was identified in a screen for ribosome associated factors (Fleischer et al., 2006); Stm1 has a variety of functions in ribosome homeostasis (Van Dyke et al., 2013); Nap1, among other functions, chaperones Rps6 and may directly facilitate 40S biogenesis (Rössler et al., 2019); Lhp1 is a yeast La protein homolog required for the maturation of tRNAs and could plausibly have a function in ribosome biogenesis (Yoo and Wolin, 1997). Pab1 is traditionally associated with mRNA processing, but also has genetic interactions with ribosome biogenesis factors (Sachs and Davis, 1990); Sro9 is an RNA-binding protein that associates with translating ribosome but also shuttles between the nucleus and cytoplasm (Röther et al., 2010); YMR310C and YGR283C are uncharacterized paralogs with predicted methyltransferase function. These ORFs, along with another nucleolar protein present in our dataset, YCR016W, are part of a large ribosome biogenesis regulon (Wade et al., 2006).

### Assignment of unmapped factors to distinct 60S intermediate structures

Pre-60S structural intermediates can be broadly divided into nucleolar, nucleoplasmic and cytoplasmic assemblies, characterized by distinct rRNA assembly states and compartment-specific RBFs. Characteristic differences in RBF composition can therefore be used to associate structurally uncharacterized RBFs with individual structures. For our analysis, we chose two representative structures for each subcellular compartment (**Figure 2A**). The first nucleolar structure is characterized by the presence of the RBFs Rrp14, Rrp15 and Ssf1/Ssf2 (Kater et al., 2017; Sanghai et al., 2018), while the second one contains the proteins Noc3, Spb1 and Spb4 (Kater et al., 2017), which engage the pre-60S after organization of rRNA domain III onto the core of the 60S and the release of Rrp14/Rrp15/Ssf1. The release of pre-60S particles from the nucleolus into the cytoplasm coincides with the organization of the L1 stalk and the incorporation of the GTPase Nog2. Our representative nucleoplasmic pre-60S particles represent states before and after rotation of the 5S rRNA, which is marked by the release of RBFs Rpf2/Rrs1 and the recruitment of the Rix1 complex (Kater et al., 2020; Wu et al., 2016). After nuclear export, final maturation occurs in the cytoplasm, as typified by the structures of early and late cytoplasmic 60S intermediates (Kargas et al., 2019; Zhou et al., 2019b). As our quantitative data mirrors these changes in RBF composition, we can use individual abundance correlations between these “marker” RBFs and unmapped RBFs to assign them to specific structural intermediates. In addition, we can expand the presence of structurally characterized factors to intermediates where they are associated with dynamic regions that are not currently resolvable in cryo-EM maps (**Figure 2A**). The correlation between early and late nucleolar RBFs and components of the 5S rRNP (uL5, uL18, Rrs1 and Rpf2) (**Figure 1C and S2**) reveals that the 5S rRNP is recruited to the maturing 60S particle at an early stage, before it is resolvable in cryo-EM reconstructions and possibly after the release of Mak11 and the binding of Nog1, Mrt4, Nsa2 and uL6. Similarly, our data indicates that the L1-stalk-interacting factors Nop2 and Nip7 as well as uL1 are already present in early nucleolar structures, although most likely in a dynamic, unmoored state compared to late nucleolar particles (**Figures 1C, 2A and S2**). During maturation, rRNAs are extensively modified by snoRNA guided enzymatic complexes, resulting in pseudouridylation and 3’-OH methylation of ribose moieties. In addition, eight large subunit rRNA bases are modified by dedicated methyltransferases (Yang et al., 2016). Our quantitative MS data, in combination with known structural intermediates, allow us to place some of these catalytic activities into the 60S timeline (**Figure 2A and S5**). Components of the box H/ACA enzymatic complex (Cbf5, Gar1 and Nhp2), while most abundant in early, structurally uncharacterized intermediates and correlating most strongly with the RBFs Nop1, Nop56 and Nop58 involved in late pre-40S processing (**Figure S5**), are also detected in the Ssf1/Rrp15/Rrp14-containing structures (**Figure 1D)**, indicating that some pseudouridylation events occur after a majority of the 60rRNA has already folded. Because the H/ACA complex is directed to methylation sites by guide snoRNA, this suggests rRNA domains III and IV, which are not ordered in these structures, are still accessible to snoRNA mediated modification. Our data also associates the activities of base-modifying methyltransferases with specific maturation intermediates: Rcm1 and Bmt2, which respectively catalyse the m^1^A and m^5^C methylation at rRNA residues 2278 and 2142, are enriched in late nucleolar particles (**Figures 1D and 2A, Figure S5**). The target nucleotides of these enzymes are located in rRNA domain IV, which is not ordered in the available nucleolar structures and therefore likely to be accessible for base modification. Another methyltransferase, Bmt5, catalyzes the m^3^U modification at nucleotide 2634. The abundance distribution of Bmt5 suggests that this modification occurs after pre-60S nucleolar release and Nog2 binding, but before 5S rRNP rotation and Rix1 binding (**Figures 1D and 2A, Figure S5**). In fact, the loop containing U2634 is solvent exposed in the pre-rotation state, but buried after 5S rRNA rearrangement (Wu et al., 2016; Zhou et al., 2019b), a process that may be promoted by the base modification.

Finally, the distribution of the C2 processing factors Las1/Grc3 correlates strongly with the binding of Rix1/Rea1, allowing us to narrowly define the timing of Las1 recruitment to the C2 site (**Figures 2A** and **S5**). This correlation between Las1 and the Rix1/Rea1 complex is consistent with the fact that Rix1-containing particles have distinct pools of particles with or without the ITS2 foot structure (Kater et al., 2020). Finally, we observe a close correlation between Ecm1 and the major export receptors Mex67/Mtr2, suggesting a functional link between these proteins in 60S nuclear export (**Figures 2A and S5**).

### Timing of DEAD-box ATPases in 60S biogenesis

Perhaps the most prominent group of proteins “missing” from current cryo-EM structures are DEAD-box ATPases. These enzymes possess ATP-dependant RNA translocation and helicase activities and play a crucial role in RNA biology and are prominently represented among ribosome biogenesis factors (Linder and Jankowsky, 2011). While the precise role of these proteins in 60S biogenesis is unknown, they are thought to facilitate ATP-driven rRNA remodelling steps during ribosome assembly (Klinge and Woolford, 2019). Of the eight DEAD-box helicases associated with 60S assembly, only Has1 has been modelled (Kater et al., 2017; Sanghai et al., 2018). However, Has1 has a structural, ATP-independent role in 60S biogenesis that is distinct from its catalytic function in 40S assembly (Dembowski et al., 2013). No high-resolution structural information exists for the remaining seven DEAD-box ATPases (Dbp6, Dbp7, Dbp9, Dbp10, Drs1, Mak5 and Spb4), although a tentative placement has been proposed for Spb4 (Kater et al., 2017; Sanghai et al., 2018), and rRNA regions contacting Mak5, Dbp10 and Spb4 have been defined by CRAC experiments (Brüning et al., 2018; Manikas et al., 2016).

In this study, all eight DEAD-box proteins are reliably identified and quantified in our dataset (**Figure 2B**), allowing us to define the timing of their engagement with pre-60S particles by analysing their quantitative distribution in our clustering analysis. Dbp6 clusters with the earliest samples (Noc1, Ras3), a placement that is consistent with previous studies showing that Dbp6 and Rsa3 form a stable complex that also includes Urb1/Urb2 and Nop8, (de la Cruz et al., 2004; Rosado et al., 2007). Dbp7, Dbp9 and Mak5, while already present in the Noc1 and Rsa3 particles, exhibit a broader distribution in our clustering pattern, extending into samples containing Ssf1 and Rrp14/15 (**Figure 2B**). Drs1 and Dbp10 are absent in the earliest clusters, but feature prominently in the intermediate nucleolar samples (**Figure 2B**). Spb4 shows up last in our timeline and uniquely extends into pre-ribosomal particles containing Rix1, suggesting that Spb4 engagement occurs last among DEAD-box proteins (**Figure 2B**). DEAD-box function appears to be restricted to the nucleolus, with only low levels observed in nucleoplasmic or cytoplasmic particles. To obtain a more detailed picture of individual pairwise interactions between DEAD-box ATPases, we calculated individual abundance distribution correlations between each ATPase and all RBFs in our pool, selecting for the strongest correlations (pairwise Pearson correlations ≥ 0.95) (**Figure 2C**; see STAR Methods for details). This analysis reveals a close abundance correlation between Dbp9, Mak5 and Mak11 and between Spb4 and Rrp17, reflecting the broad pattern discussed above and suggesting a role for Dbp9 and Mak5 in the early stages of nucleolar assembly and a late nucleolar role for Rrp17(Oeffinger et al., 2009).

### Implementation of a mono- and intralink filter (mi-filter) to improve false-discovery rates in AP-MS XL-MS data

Obtaining an overall timeline for 60S biogenesis using our AP-MS derived label-free quantification data can be achieved even with significant “noise” contributions, given the typically low false-discovery rates for these experiments (Hein et al., 2015; Tyanova et al., 2016a; Tyanova et al., 2016b). However, potentially higher false discovery rates can undermine the confidence in individual protein-protein interactions (PPIs). Current state-of-the-art XL-MS workflows are challenged by low-abundance proteins (Fursch et al., 2020), and their false discovery rates (FDRs) are frequently underestimated (Beveridge et al., 2020). Because the localization of previously unmapped or novel RBFs relies entirely on inter-protein crosslinks (interlinks), the high FDRs often associated with interlinks represent a challenge to our affinity enriched XL-MS dataset (Erzberger et al., 2014).

To increase the reliability of our dataset and to enable the localization of completely novel or previously unmapped RBFs with high confidence, we set out to filter out false-positive PPIs. To date, FDR assessment in XL-MS has been primarily addressed through the optimization of scoring algorithms and the use of decoy databases (Beveridge et al., 2020; Fischer and Rappsilber, 2017; Gotze et al., 2019; Liu et al., 2015; Walzthoeni et al., 2012). Here, we took a different approach and used the expected distribution of monolinks, intralinks and interlinks in our sample to filter out false positive PPIs: if a protein is present at high-enough quantities to be detectable by XL-MS, monolink and intra-protein crosslinks will be generated at higher rates than interlinks (monolinks are formed when only one of the two the active groups of the crosslinker is able to react with a lysine side-chain) (Fursch et al., 2020). The mi-filter in our workflow stipulates that only proteins identified with at least one monolink or intra-protein crosslink within our dataset can be considered a bona fide interlink representing a legitimate PPI. This simple and intuitive filtering has a dramatic effect on our analysis: Using the Rix1 dataset as an example, a “standard” filter setting relying on one identified high-confidence crosslink per unique crosslinking site results in high FDRs, in particularly for inter-protein crosslinks (**Figure 3 A, D, G, J**). A more stringent filter setting relying on two independently identified high-confidence crosslinks improves the overall FDR to ∼4 %, a 7.5 % reduction, but still results in a high percentage of unreliable interlinks and an overall low degree of confidence in individual PPIs (**Figures 3 B, E, H, K**). Application of the mi-filter however results in a dramatically improved overall FDR of <0.5 %, and most strikingly, yields a near 100-fold improvement in the FDR for inter-protein crosslinks (**Figure 3 C, F, I, L)**. By applying the mi-filter to our complete crosslink dataset, the interlink FDR is improved to <0.4 %, ensuring that the filtered PPIs can be mapped with great confidence (**Figure 3 M, N, O)**.

**Figure 3:**
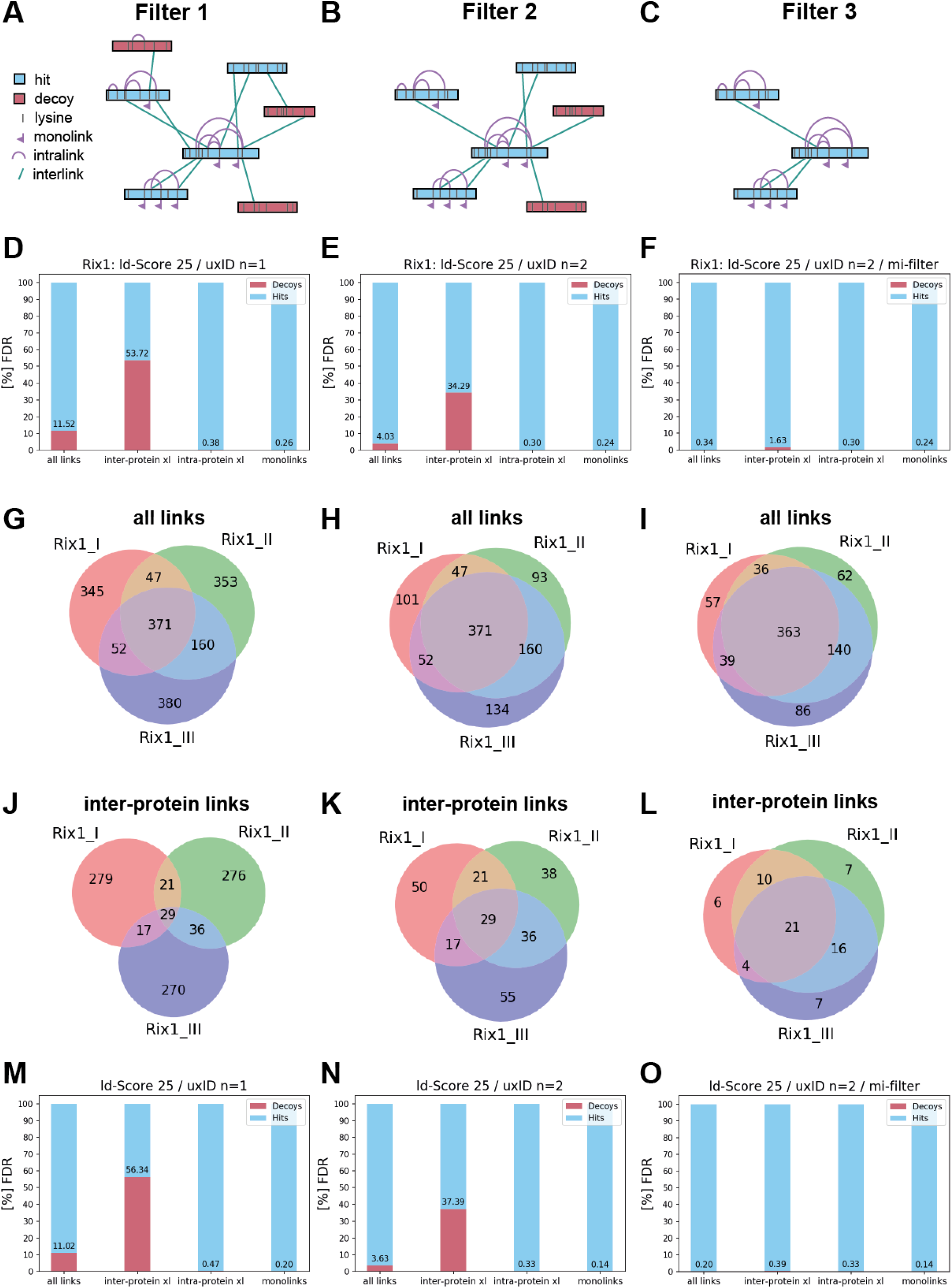
Mono- and intralink filter (mi-filter) for improved false-discovery rates in large scale AP-MS XL-MS data. Illustration of the three different filter criteria used to test their influence on the false-discovery rate (FDR) in our crosslink dataset. Shown are true positive crosslinks (hits) in blue and false-positive (decoys) in red. (**A**) The “standard” filter setting relies on one identified high-confidence crosslink per unique crosslinking site. (**B**) The more stringent filter setting relies on two independently identified high-confidence crosslink per unique crosslinking site (among all datasets). (**C**) For the final filter setting the mono- and intralink filter (mi-filter) was applied. The mi-filter requires that for each protein involved in an inter-protein crosslink also at least one mono- or intralink had to be detected in one of the biological replicates. (**D, E, F**) Bar charts for each filter setting and the respective FDRs (shown in percent) for all link types for the Rix1 dataset. (**G, H, I**) Venn diagrams showing the overlap between the three independent biological replicates of the Rix1 particle for all link-types and (**J**, **K**, **L**) for inter-protein crosslinks only. **(M**, **N**, **O**) Bar charts for each filter setting and the respective FDRs (shown in percent) for all link types in our whole dataset. Please note that for H, I, K and L all links, which were identified in only 1 biological replicate of Rix1 pre-ribosomal particles, were also identified in at least one other pre-ribosomal particle.

### A database of high-confidence interlinks in 60S biogenesis intermediates

Our final crosslink dataset, after application of the mi-filter, consists of over 60000 mono- and crosslinks, with 2844 inter-protein crosslinks among 145 individual proteins (**Supplementary Data 2**). 71 of these involve r-proteins of the large or small subunit (including homologous r-proteins) and another 50 are associated with structurally characterized RBFs (46 60S factors and 4 90S/40S factors). Only 7 RBFs known to be present in pre-60 intermediates are not represented by an interlink in our database (Tif6, Bud20, Sdo1, Ria1, Reh1, Mak16, and Sdo1). The remaining 352 interlinks involve 24 proteins with currently unknown localization (**Supplementary Data 2**). In order to visualize the sites of action of these remaining RBFs, we overlaid crosslink networks on the outlines of cryo-EM structures of pre-ribosomal particles, choosing the most appropriate structural scaffold, although some datasets, notably Cic1, span diverse structural intermediates (**Figures 2A and 4, Figure S7)**; for an overview of the highlighted regions see **Figure S6**). The 22 RBFs with previously unknown localization as well as 2 candidate biogenesis factors are shown together in **Figure 5A**.

**Figure 4:**
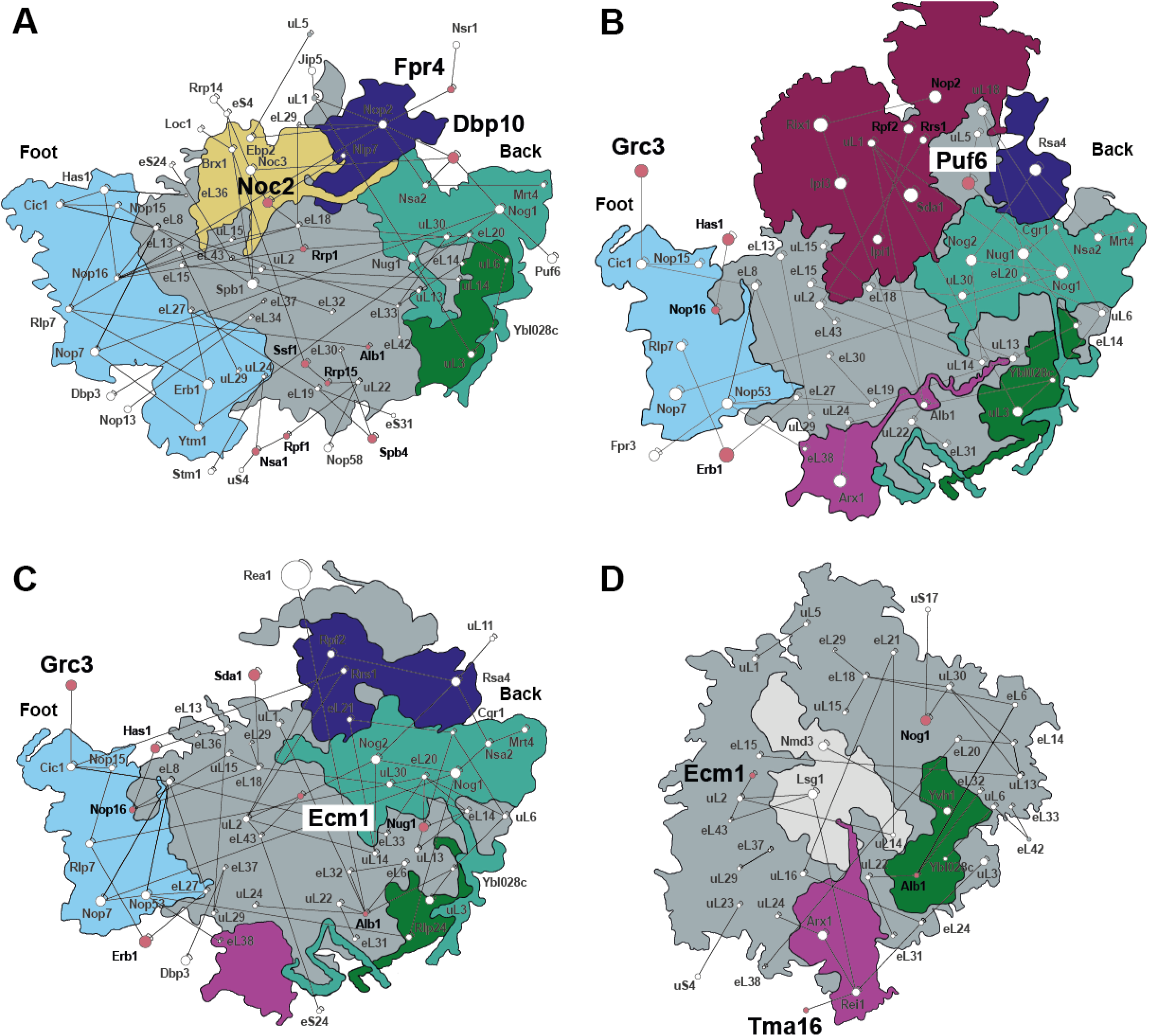
High-confidence crosslink networks for select RBFs mapped onto corresponding structures of pre-ribosomal particles. Shown are high-confidence crosslinks using our mi-filter for a select subset of RBFs from our AP-MS XL-MS dataset mapped onto corresponding previously published cryo-EM reconstructions. (**A**) Crosslinks of the Ytm1 pre-ribosomal particle mapped onto pdb 6ELZ. (**B**) Crosslinks of the Rix1 pre-ribosomal particle mapped onto pdb 6YLH. (**C**) Crosslinks of the Nog2 pre-ribosomal particle mapped onto pdb structure 3JCT. (**D**) Crosslinks of the Lsg1 pre-ribosomal particle mapped onto pdb 6RZZ. RBFs that were identified within a particle-specific crosslinking dataset (n=2) but are not present in the respective PDB structure (but are present in some other PDB structure) are indicated in red and labelled bold, while RBFs and factors with presently unknown position (Noc2, Fpr4, Dbp10, Grc3, Puf6, Ecm1 and Tma16) are labelled with a larger font size. For a description of the highlighted regions see Figure S6. Please note that proteins containing only intra- or monolinks are not shown.

**Figure 5:**
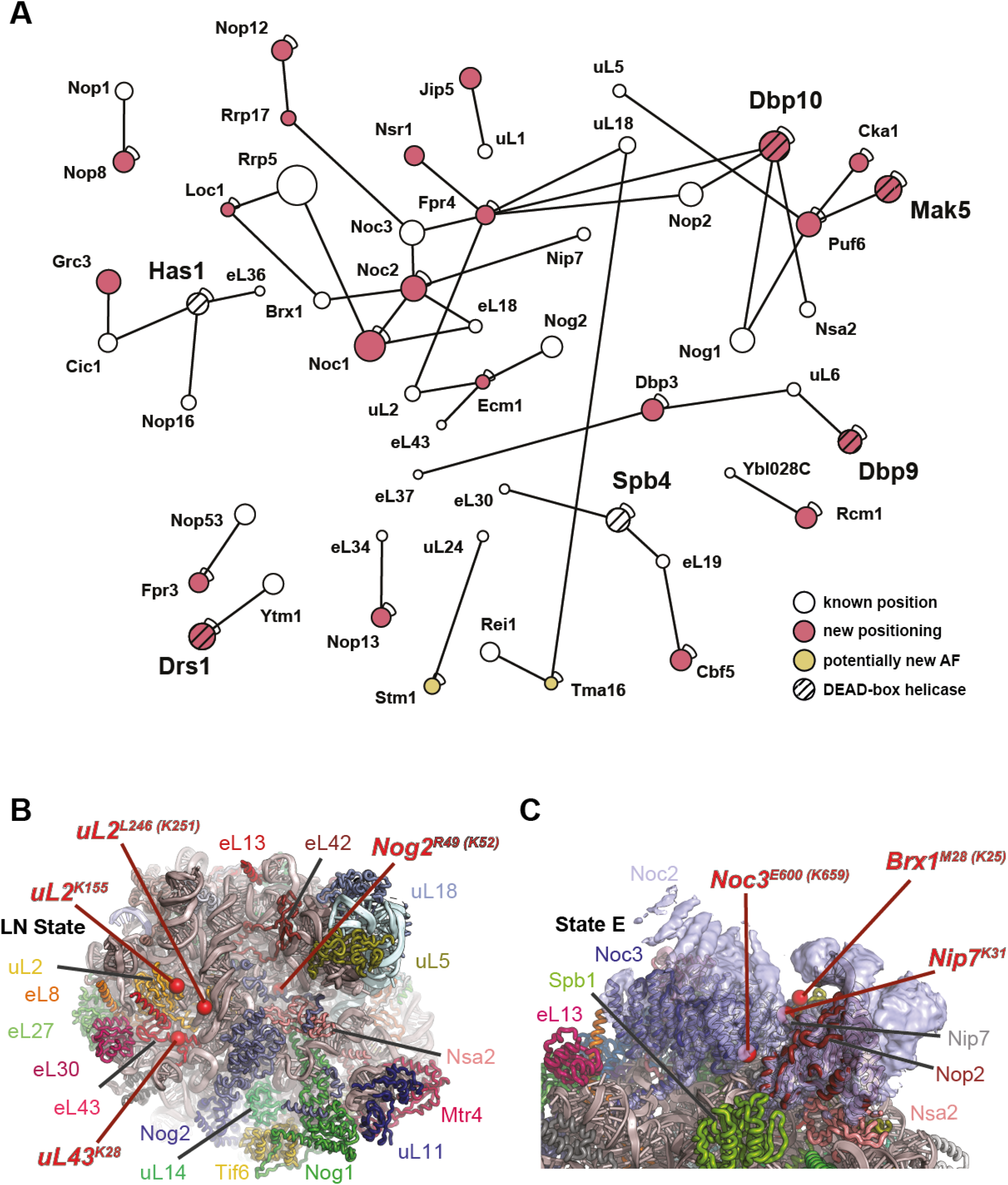
Placement of structurally uncharacterized RBFs. (**A**) Map of mi-filtered high-confidence crosslinks for candidate novel biogenesis factors and all RBFs whose localization within pre-ribosomal particles are currently not known from our complete AP-MS XL-MS dataset. New localizations for known RBFs are shown in red and for novel RBFs in yellow. DEAD-box helicases additionally contain black stripes. (**B**) Placement of Ecm1 on nucleoplasmic pre-60S particles. Cartoon representation of pdb 6N8J with key proteins labeled. Residues with interlinks (or closest modeled residue if disordered) to Ecm1 are shown as red spheres and labeled. The only access to residue K52 of Nog is in the post-rotation state before Nmd3 binding. (**C**) Assignment of Noc2 to uncharacterized EM density in late nucleolar 60S particles. Cartoon representation of pdb 6ELZ with overlayed EM density (EMDB 3891), low pass filtered to 4 Å. Key proteins are labeled. Red spheres mark the position of residues (or closest modeled residue if disordered) of Noc3, Brx1 or Nip7 that form interprotein crosslinks to Noc2. The features of the density are consistent with the prediction that Noc2 forms helical repeats and directly interacts with Noc3.

### Identification and localization of Tma16, Las1/Grc3, Ecm1 and Noc2

While our database will undoubtedly aid future interpretation of cryo-EM maps of 60S structural intermediates, specific interactions identified in our interlinks can already be used to map certain RBFs to 60S structural intermediates. Of the nine candidate novel biogenesis factors identified in our quantitative analysis, only two, Tma16 and Stm1, are also represented in our inter-protein crosslink library. In total, 5 inter-protein crosslinks between the N-terminal part of Tma16 and Rei1 and 5 interlinks between the C-terminal region of Tma16 and uL18 were identified (**Supplementary Data 2**) in pre-ribosomal particles purified with affinity tagged Arx1 (**Figure S7H**) and Lsg1 (**Figure 4D**). In our quantitative MS analysis, Tma16 exhibits an abundance pattern most similar to the Arx1/Alb1 complex (data not shown) across the Cic1, Rix1, Nop12, Nog2, Arx1 and Lsg1 pulldowns (**Figure S4**). Together, this data points to Tma16 as a novel pre-60S cytoplasmic RBF. uL18 and Rei1 are located quite distantly from each other on the surface of the pre-60S and engage opposite termini of Tma16, suggesting an extended conformation for this factor on the 60S surface.

During ribosome biogenesis, ITS2 removal is initiated by Las1-mediated cleavage at the C2 site within ITS2, followed by phosphorylation of the resulting 5’-hydroxyl product on the 25S rRNA by the Grc3 polynucleotide kinase (Fromm et al., 2017; Gasse et al., 2015; Pilion et al., 2019). The activities of Las1 and Grc3 are coupled within the tetrameric assembly (Pilion et al., 2019). In our interlink dataset, we map an interaction between Grc3 and the structurally uncharacterized C-terminal domain of Cic1 in pre-ribosomal particles purified with affinity-tagged Rix1 (**Figure 4B**) and Nog2 (**Figure 4C**). Pearson correlation analysis based on protein abundance in the different pre-ribosomal particles shows that the Las1/Grc3 complex correlates strongly with the Rix1/Rea1 complex (**Figure S5**), in line with the structural observation that pre-60S particles containing Rix1 and Rea1 can be reconstructed both with and without a foot structure (Kater et al., 2020). The correlated recruitment of the Rix1/Rea1 complex and Las1/Grc3 hints at a possible coordination between Rix1 complex binding and ITS2 processing.

Nuclear export of pre-60S particles is mediated by the export factors Arx1, Mex67-Mtr2, Nmd3 and Ecm1. These factors have overlapping, non-redundant roles in pre-60S export (Yao et al., 2010). While structural information is available for the other export factors, Ecm1, which binds to nucleoporins and shuttles between the nucleus and the cytoplasm, has not been observed in any cryo-EM reconstructions to date. Ecm1 is abundantly featured in our interlink dataset, with a total of 68 unique inter-protein crosslinks between Ecm1, uL2 and eL43 found in the Nop12, Nog2, Arx1 and Lsg1 datasets and 3 crosslinks to Nog2 exclusively observed in the Nog2 dataset (**Supplementary Data 2**) (**Figure 4, 5 and S7**). Mapping of these interacting residues onto late nuclear pre-60S particles implies that Ecm1 binds 60S after the removal of Rsa4 by Rea1, but before the release of the GTPase Nog2, because this is the only structural intermediate where Nog2 is accessible for crosslinking with Ecm1 (**Figure 5B** (Zhou et al., 2019b). In our MS quantification analysis, Ecm1 correlates strongly with Mex67 and Mtr2 and exhibits a similar abundance pattern within late nucleoplasmic and cytoplasmic intermediates (**Figure S5**), implying a concerted engagement of pre-60S particles by these export factors.

Noc2 is an unusual pre-60S RBF, forming two distinct complexes (Noc1/Noc2 and Noc3/Noc2) that act at different stages of maturation. Noc3 and Noc1 share a conserved Noc motif that likely mediates their interaction with Noc2 (Milkereit et al., 2001). Accordingly, while the abundance of Noc1 and Noc3 is limited to very early and late nucleolar intermediates, respectively, Noc2 shows a bimodal distribution that matches the presence of Noc1 and Noc3, but not the intervening pre-60S intermediates (**Figure 1C** and **1D**). Independently, we detect 10 interlinks between Noc1 and Noc2, all from a particle pool purified with tagged Noc1 (**Figure S7, Supplementary Data 2**). The Noc3/Noc2 interaction is represented by 67 independent crosslinks, all mapping to late nucleolar samples. This pattern is consistent with two independent pre-60S engagements by Noc2, mediated by distinct dimerization partners. In late nucleolar samples, Noc2 is also associated with Brx1 (4 interlinks) and Nip7 (12 interlinks). Mapping these residues (or their closest modelled residue neighbours) onto the structure of late nucleolar 60S substrates shows that they triangulate around an area of unassigned density above Noc3 (**Figure 5C**). This density is consistent with the helical repeats predicted to occur within Noc2 (Milkereit et al., 2001). Because all crosslinks involve residues within the last 110 residues of Noc2, we propose that the Noc2 helical repeats extend away from Noc3 in a C- to N-terminal direction.

### The Ck2 complex engages early 60S biogenesis intermediates

The heterotetrameric casein kinase 2 (Ck2) complex has a central role in cell growth and proliferation, acting as a serine/threonine kinase on a diverse set of protein targets (Kos-Braun et al., 2017). Ck2 is composed of two catalytic alpha subunits (Cka1 and Cka2) and two regulatory subunits (Ckb1 and Ckb2). Ck2 is a component of the CURI complex, along with the small subunit RBFs Rrp7, Utp22 and Ihf1, and has an established role in 40S biogenesis (Krogan et al., 2004). More recently, Ck2 activity was found to be essential to trigger a switch in ITS1 processing during stress responses (Kos-Braun et al., 2017). Our data suggests that Ck2 involvement in ribosome biogenesis extends to the large subunit as well. All 4 subunits of Ck2 show a similar protein abundance pattern in ‘early’ and ‘intermediate’ pre-ribosomal particles purified with affinity-tagged Ssf1, Nsa1, Nop16, Ytm1, Cic1 and Noc1 (**Figure 6A**). In addition, we find two high-confidence inter-protein crosslinks between Cka1 (CSK21) and Puf6 in both Noc1 and Nop16 samples. To confirm the link between Ck2 and the large subunit, we carried out an additional, reciprocal pulldown using TAP-tagged Cka1 (CSK21). In this pulldown, we identified an inter-protein crosslink between Cka1 and uL13 together with mono- and intralinks among various proteins of the pre-60S foot structure and within Ssf1/Rrp15/Rrp14, Brx1/Ebp2 and Mak16/Rpf1/Nsa1 which are all present in the early nucleolar pre-60S structures (**Supplementary Data 2**). Correlations from our quantitative MS data (**Figure 6B**) as well as interlinks and known physical interactions allow us to generate an extensive interaction map involving Ck2 (**Figure 6C**). This interactome identifies Ck2 as a bona-fide component of early pre-60S particles with an important role in licensing the earliest assembly steps in 60S maturation. A number of unmapped factors, such as Puf6/Loc1, Fpr3/4 and the DEAD-box helicases Drs1, Mak5 and Dbp9 are connected to Ck2 and may be linked to its function during early nucleolar 60S maturation (**Supplementary Data 2**).

**Figure 6:**
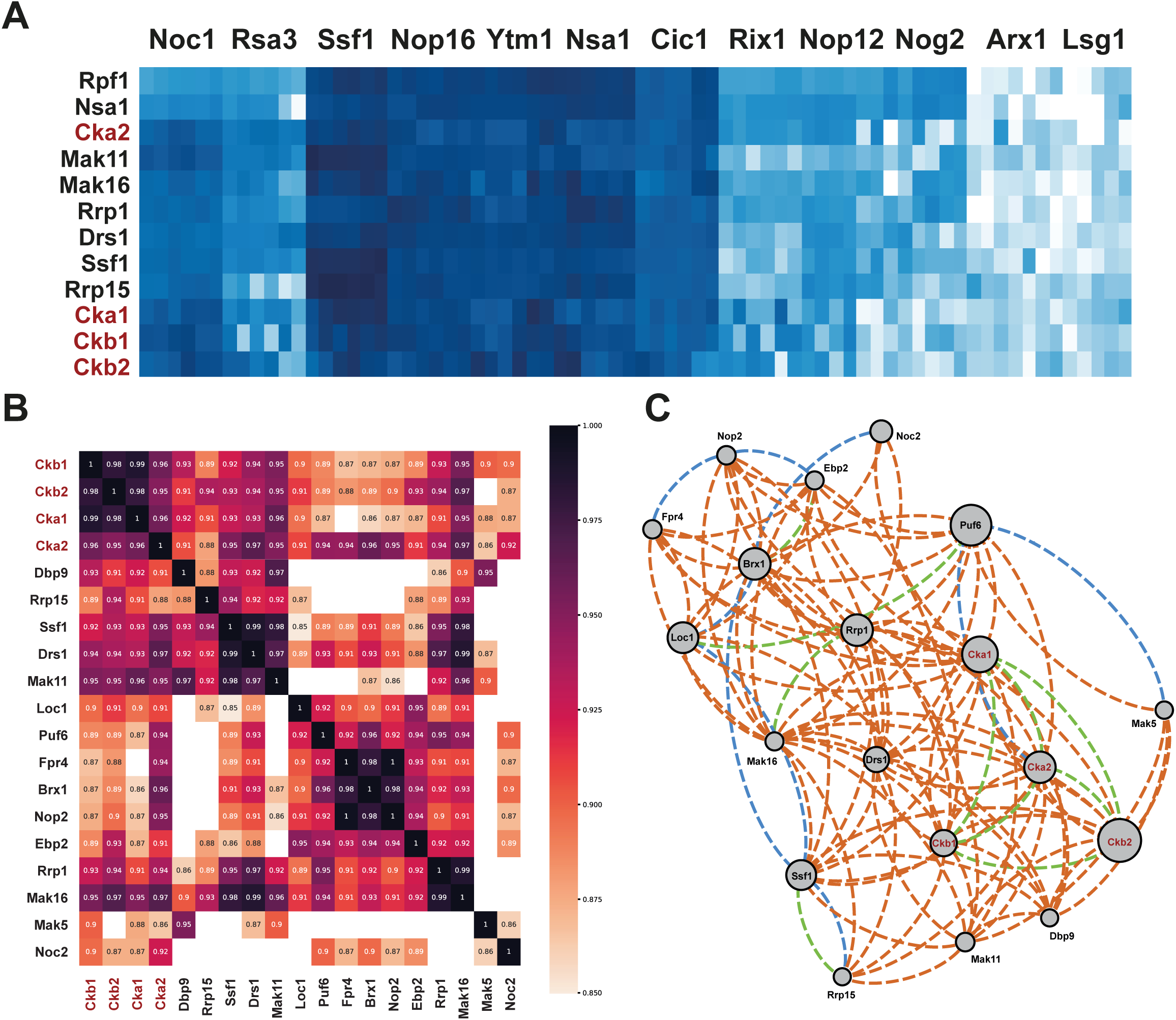
The casein kinase complex is a 60S biogenesis factor. (**A**) Protein abundance subcluster for the Ck2 complex (consisting of Cka1, Cka2, Ckb1 and Ckb2) from our overall AP-MS data by hierarchical cluster analysis of the ‘Intermediate’ proteins (Figure 1, blue cluster) using an L1 distance metric. Protein abundances are shown as Z-score normalized log2 transformed LFQ values ranging from ≤ −4.0 (white) to 0.0 (azure blue) to ≥ 2.5 (dark blue). (**B**) Heatmap of pairwise correlations within the Ck2 interactome. To be included, factors had to have a correlation of >0.85 with three out of the four subunits of the Ck2 complex, one of which had to be ≥ 0.9. (**C**) The Ck2 interactome during 60S biogenesis. Shown are interactions among proteins with correlations from Figure 6B (orange lines), high-confidence interlinks (blue lines) and direct physical interactions (green lines). These interactions are thought to occur during the earliest stages of nucleolar 60S maturation.

### Localization of the DEAD-box helicases Dbp9 and Dbp10

Except for Dbp6 and Dbp7, all of the DEAD-box helicases involved in 60S biogenesis and identified in our MS quantification are represented in our crosslinking dataset (**Figure 5A**). Dbp9, Drs1 and Mak5 are contained in the interactome around Ck2, forming interlinks with uL6, Ytm1 and Puf6, respectively (**Figure 6**). Drs1 engages the Ytm1 β-propeller while it is still loosely tethered to the pre-60S core, making precise mapping of Drs1 impossible. Similarly, Mak5 is linked to Puf6, which has not been identified in any cryo-EM maps. Dbp9, on the other hand, forms a link between a loop insertion in its N-terminal RecA domain and uL6. Structures of nucleolar intermediates showed that uL6 binds the pre-60S core after domain VI docks onto the core (**Figure 7A**) (Kater et al., 2017; Sanghai et al., 2018). The intermediate immediately preceding uL6 binding is characterized by the presence of Mak11, while uL6 is necessary for the binding of the Nog1 GTPase domain, Mrt4 and Nsa2. The timing of uL6 binding and the absence of Dbp9 in later nucleolar particles suggests that Dbp9 performs its catalytic function during a narrow window in time: after Dbp9 binding, but before release of RBFs Ssf1, Rrp14 and Rrp15. In fact, the relative inaccessibility of the uL6^K54^ in later intermediates suggests that the function of Dbp9 may be directly linked to the rRNA rearrangements necessary for Nog1-GTPase domain, Mrt4 and Nsa2 recruitment.

**Figure 7.**
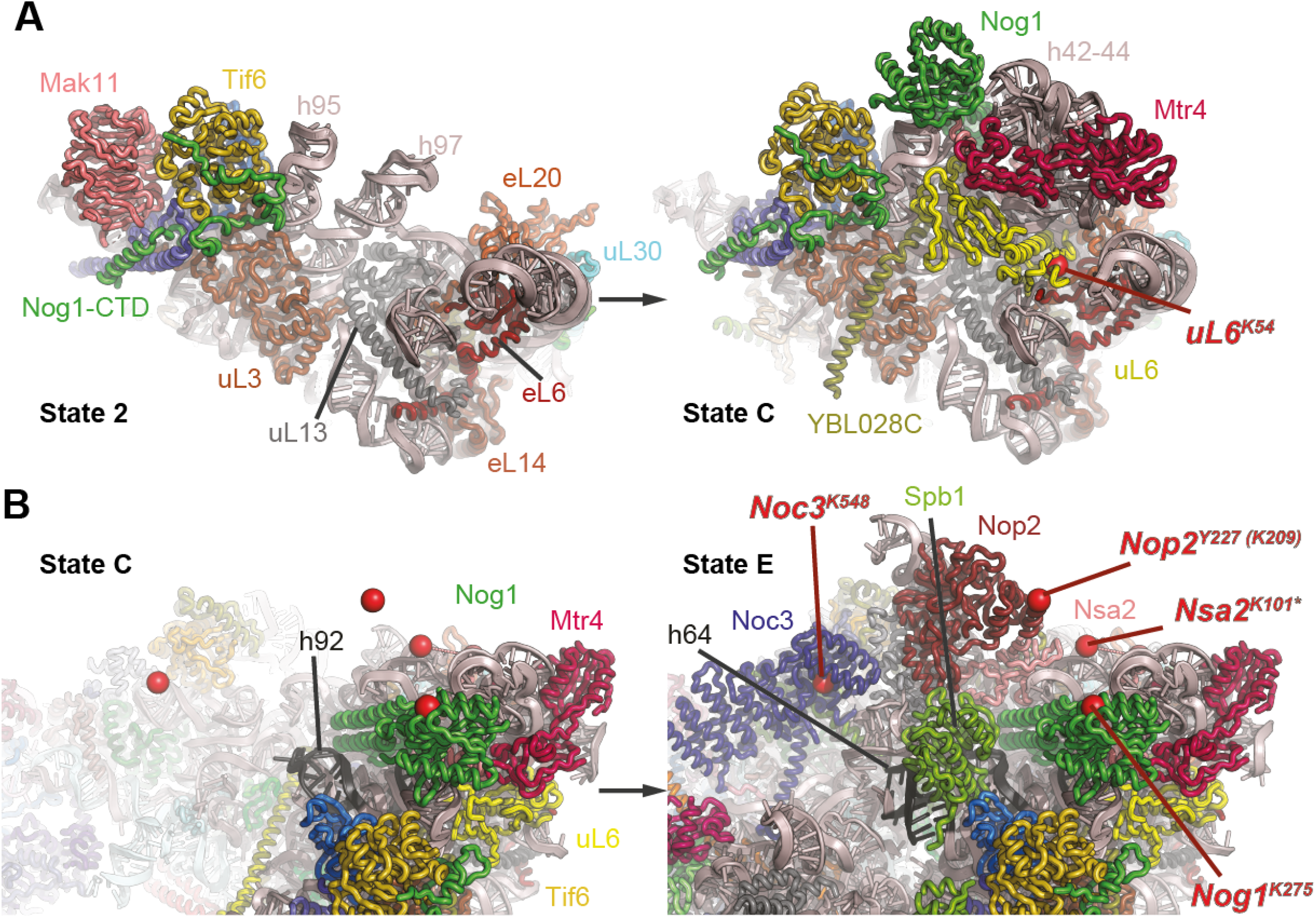
Placement of DEAD-box helicases Dbp9 and Dbp10. (**A**) Cartoon representation of early nucleolar pre-60S particles (left – 6C0F, right – 6EM1), showing the final assembly of RBFs engaging rRNA domain VI. The transition is defined by the release of Mak11 and the binding of uL6, Nsa2 and Mrt4, as well as the N-terminal GTPase domain of Nog1. The N-terminal RecA domain of Dbp9 crosslinks to uL6, indicating that it engages 60S during the final assembly of this region of the 60S ribosome. (**B**) Dbp10 engages the pre-60S ribosome after the Noc3 binding, but before engagement of helix 92 by the methyltransferase domain of Spb1. LEFT – cartoon depiction of nucleolar pre-60S intermediate (pdb 6EM1) with key proteins labeled. Residues with specific crosslinks to Dbp10 are shown as red spheres, revealing that the majority of interacting residues are not yet ordered. RIGHT - Crosslinked residues are all present in the late nucleolar 60S structure (pdb 6ELZ), but the likely interaction site with rRNA on helices h64 and h92 is occupied by the methyltransferase domain of Spb1, suggesting that Dbp10 engages the pre-60 before Spb1 catalysis.

Our interlink database also provides extensive positional information about the DEAD-box proteins that engage late nucleolar 60S intermediates, Dbp10 and Spb4. Our Spb4 crosslinks confirm the assignment of Spb4 to a distinct, bilobed feature in the late nucleolar cryo-EM maps next to eL19, eL30 and Ytm1 (Kater et al., 2020; Sanghai et al., 2018). Spb4 binding is preceded in our timeline by Dbp10. A total of 65 interlinks associate Dbp10 with the RBFs Nog1 (3 links), Nop2 (2 links), Nsa2 (47 links) and Noc3 (13 links). On Dbp10, the residues proximal to Nog1, Nop2 and Nsa2 are located in the C-terminal extension of Dbp10, suggesting that this segment binds pre-60S particles in the cleft between rRNA domain VI and the L1 stalk (**Figure 7B**). The link to Noc3 is located within the N-terminal RecA domain, placing the enzymatic core of the protein near the binding site of the N-terminal methyltransferase domain of Spb1 (Spb1-MT). CRAC experiments identified rRNA helices h89-h92 and h64 as Dbp10 interacting regions (Manikas et al., 2016). In the structure of a late nucleolar 60S intermediate, these rRNA elements are engaged by the MT domain of Spb1 and are not accessible for Dbp10 binding (Kater et al., 2017). Taken together, this suggests that Dbp10 binds the 60S prior to Spb1-MT and Spb4 binding, in line with our quantitative MS data. In fact, by engaging helices h89-h92 and h64, Dbp10 blocks these rRNA elements from being available to bind the Spb1-MT domain. Further studies will be needed to define the mechanistic details of the sequential engagement of these RNA elements by Dbp10 and Spb1-MT.

## Discussion

Understanding the assembly process of large molecular complexes remains a challenging problem, especially the molecular characterization of transient and dynamic intermediates. The integration of both quantitative and structural proteomics data with atomic resolution models is a powerful approach to address these gaps. Applying this strategy to large subunit biogenesis, we selected twelve 60S RBFs encompassing the full maturation cycle as bait proteins to generate a comprehensive timeline for nearly all known RBFs, including 9 newly identified candidate RBFs. In order to correlate the information from our quantitative proteomics data with known structural intermediates, reliable information of PPIs based on inter-protein crosslinks is absolutely essential. Our simple and intuitive mi-filter, which removes interlinks if the linked polypeptides are not additionally represented within their respective intralink or monolink pools, improves interlink FDRs more than a 100-fold and therefore significantly enhances our ability of each and any individual interlink to provide reliable positional information for the molecular description of transient intermediates. Our resulting high-confidence interlink dataset reliably recapitulated the positioning information for structurally characterized 60S RBFs and reveals novel positioning information for 22 known, but currently structurally uncharacterized 60S assembly factors. In addition, two of our 9 candidate novel RBFs, Tma16 and Stm1, are also represented in our interlink dataset, further suggesting a role for these factors in cytoplasmic 60S maturation.

Combining our quantitative and positional MS data with available cryo-EM structures of pre-60 ribosomal intermediates allowed us to precisely map interaction areas for the RBFs Noc2, Las1/Grc3 and Ecm1 during key steps of 60S maturation: Interlinks to Noc2 associate it unambiguously with a series of helical repeats above Noc3 in late nucleolar structures. Our analysis shows that the ITS2 processing complex Las1/Grc3 is recruited to early nucleoplasmic 60S particles at the same time as the Rix1/Rea1 complex, suggesting a coordination between ITS2 removal, 5S rRNA rotation and Rsa4 removal. Ecm1, one of several 60S export factors, is closely linked with the presence of the major export receptor, Mex67/Mtr2, suggesting that the assembly of an export-competent 60S particle occurs in a concerted manner.

Our data on the heterotetrameric Ck2 casein kinase complex reveals an extensive interaction network between this regulatory complex and a cluster of nucleolar RBFs, extending the role of Ck2 to 60S maturation and providing a framework to understand the function of this multifunctional kinase in ribosome biogenesis. Ck2 is an essential component of TOR-mediated stress-responses that leads to an alternate processing of ITS1 and stalling of 60S biogenesis. To date, the precise signalling mechanism between TOR and Ck2 has not been elucidated, but systematic studies of Ck2 substrates in mammalian cells identify the human homologs of two of our proposed 60S Ck2 interaction partners, Nop2 and Rrp1, as potential Ck2 phosphorylation targets (Rusin et al., 2017). Importantly, our data, including the purification of bona fide 60S intermediates with affinity-tagged Cka1, suggests that it is not just the catalytic activity of Ck2 that is essential for its role in maintaining biogenesis, but that Ck2 is physically associated with a subset of nucleolar ribosome intermediates during early 60S maturation.

To date, the most elusive RBFs in 60S biogenesis have been the seven DEAD-box ATPases whose catalytic activity is essential for nucleolar 60S maturation. We identify all of these helicases in our quantitative dataset and all but Dbp6 and Dbp7 in our crosslinking analysis, defining a timeline of 60S engagement for these proteins: Dbp6 à Dbp7 à Mak5 à Dbp9 à Drs1 à Dbp10 à Has1 à Spb4. Dbp6, which acts within a complex composed of the factors Urb1, Urb2, Rsa3 and Nop8, engages the emerging 60S particle just as ITS1 is being processed and the 35S separated into the 20S and 27S fragments. Mak5 and Dbp9 are part of our Ck2 interaction cluster and likely play a role in organizing the docking of rRNA domain VI to the 60S core. Because of its direct interaction with uL6 and the fact that the timing of uL6 incorporation into the pre-60S is known from cryo-EM structures of 60S intermediates, we propose a distinctively timed function for Dbp9 in the assembly of the Nog1/Nsa2/Mrt4 region of nucleolar pre-60S intermediates. Similarly, an extensive network of interlinks defines the placement of Dbp10, and establishes that the binding of Dbp10 and that of the methyltransferase domain of Spb1 must occur sequentially. Both Dbp10 and Spb4 are associated with rRNA domain IV, the last segment of the rRNA to dock against the pre-60S core, and may remodel this rRNA region to chaperone its folding and ordered assembly.

In summary, in this work we successfully applied quantitative proteomics and XL-MS together with large scale biochemical enrichment of pre-ribosomal particles as a stand-alone technique to characterize transient and dynamic interactions across the full landscape of 60S pre-ribosomal particles. Integration of our MS data with available cryo-EM structures allowed us to comprehensively map RBFs involved in 60S ribosome biogenesis, providing important new insights into the timeline of Ck2 involvement and DEAD-box ATPase function in 60S biogenesis. Our high-confidence large-scale crosslinking interaction map and detailed timeline represents an essential resource for future structural and functional studies of 60S ribosome biogenesis.

## Acknowledgements

We thank Christoph Hanselka for help with reviewing of the crosslink database, Antonia Vogel, Niginia Borlinghaus and Philipp Schmid for help with the affinity enrichments and Kai-Michael Kammer and Robin Marzucca for help with python/ pandas scripts for FDR calculation and data analysis. We thank the lab of Elke Deuerling for *S. cerevisiae* strains Arx1-TAP, Lsg1-TAP and wild type BY4741. This work was supported by the German Research Foundation Collaborative Research Centre 969 (Project A06). J.P.E. was supported by the Cancer Prevention and Research Institute of Texas (RR150074), the Welch Foundation (I-1897), the UTSW Endowed Scholars Fund and the National Institutes of Health (GM135617-01). F.S. is grateful for funding from the Emmy Noether Programme of the German Research Foundation (STE 2517/1-1).

## Author Contributions

C.S., J.P.E. and F.S. conceived the study and experimental approach. C.S. expressed, purified and crosslinked pre-ribosomal particles with help from J.J.. C.S. implemented and applied the mi-filter. C.S., J.J., J.E. and F.S. analysed the data. C.S., J.P.E. and F.S. wrote the paper with input from all authors

## STAR methods

### Key Resources Table

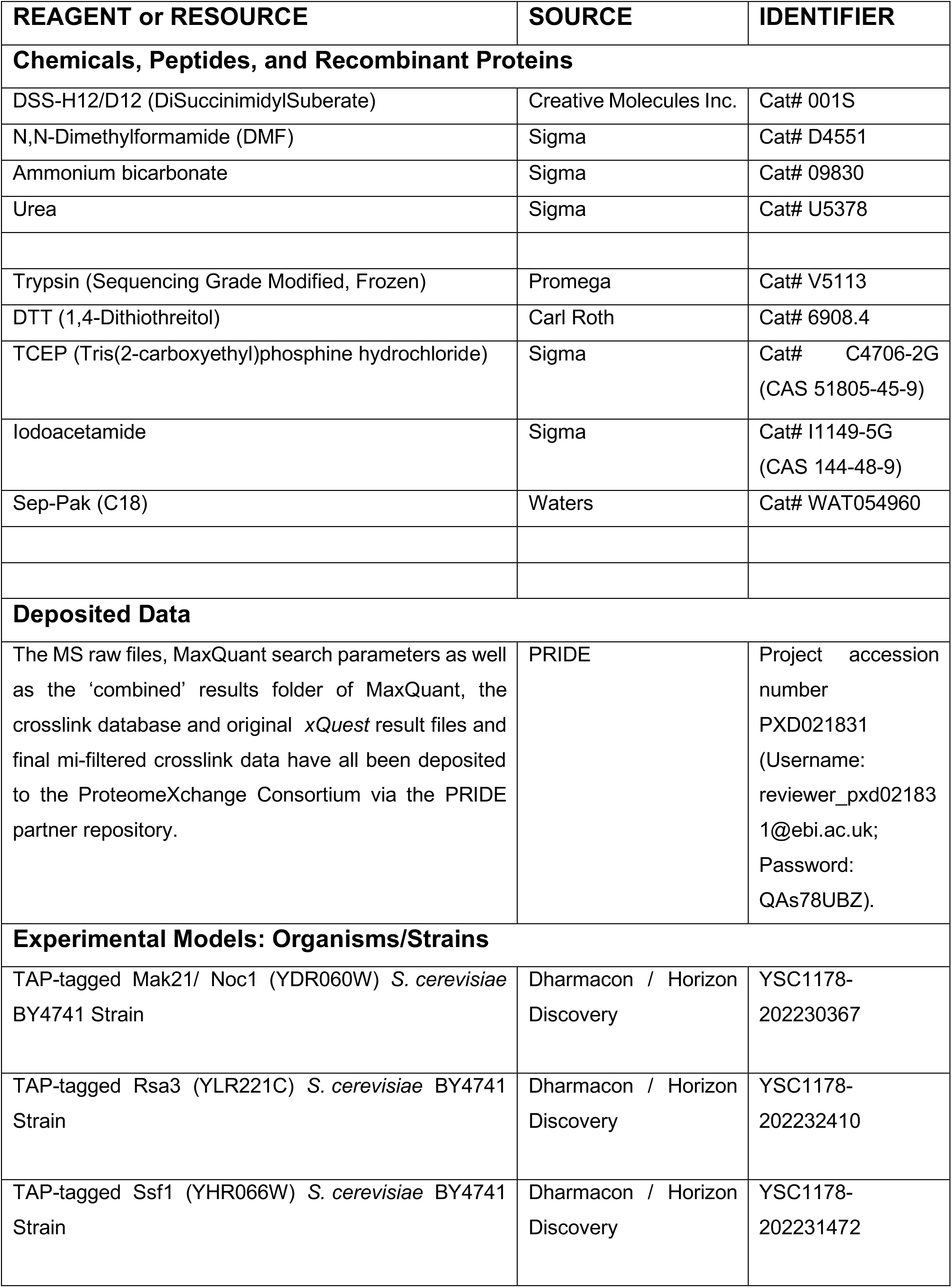
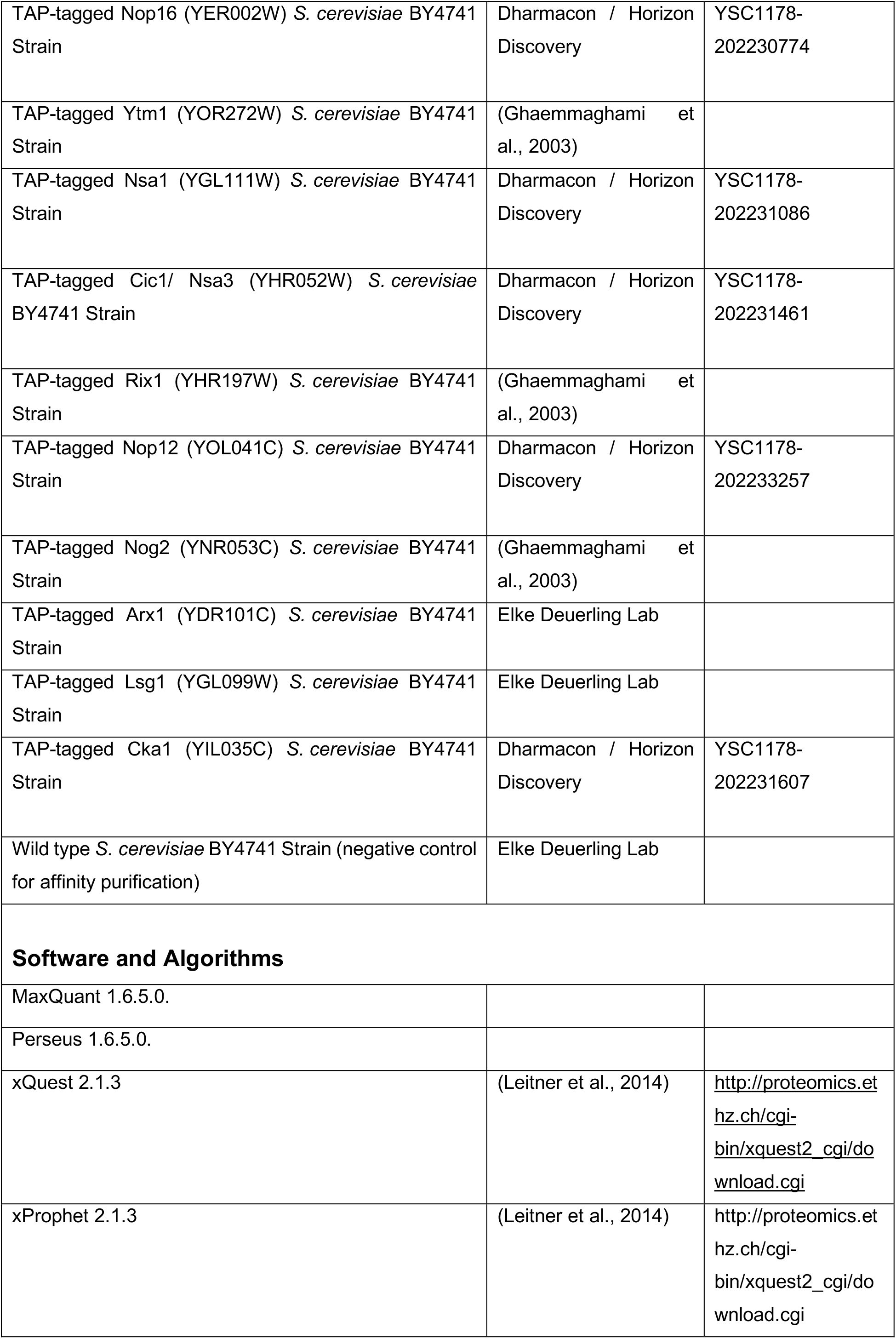
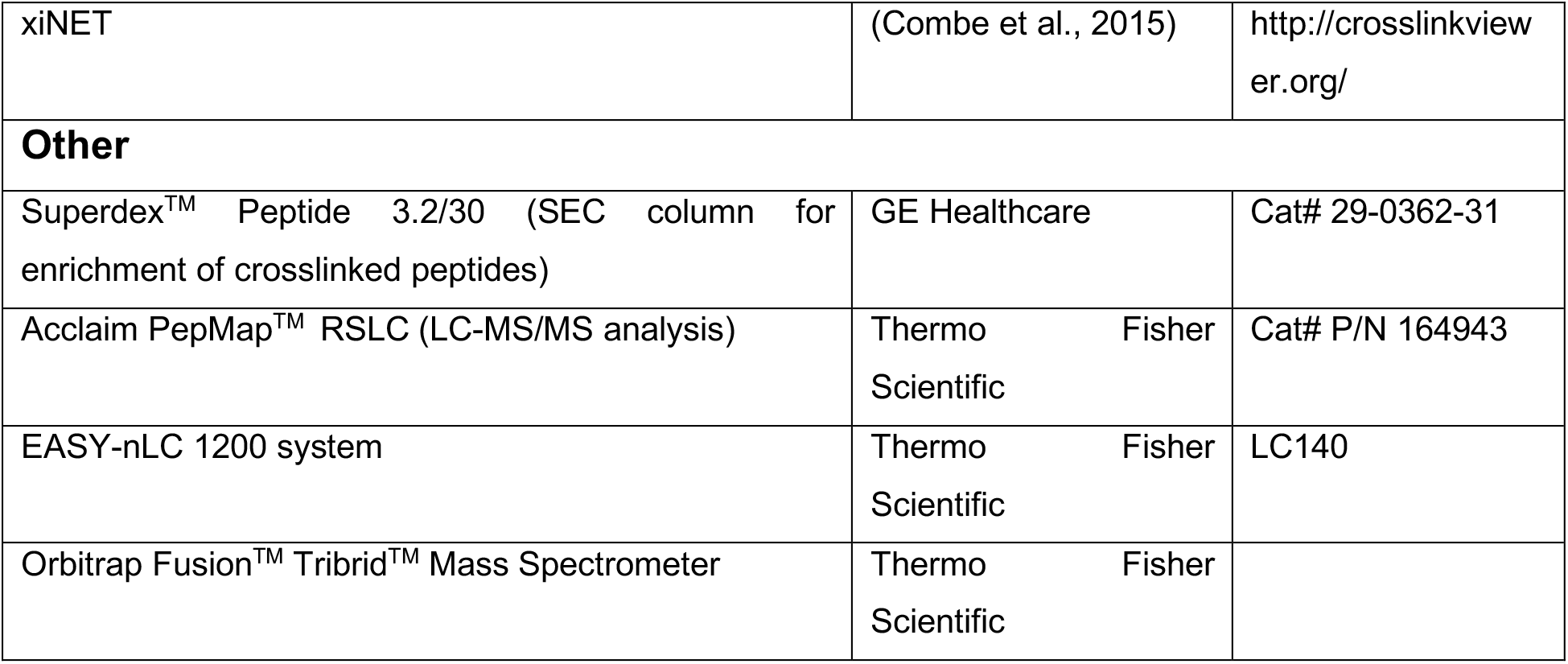

### Affinity purification and chemical crosslinking of ribosome assembly intermediates

Ribosome assembly intermediates were purified from *S. cerevisiae* BY4741 cells by tandem affinity purification (TAP). 12 different *S. cerevisiae* BY4741 strains with the TAP-tagged bait proteins Noc1, Rsa3, Ssf1, Nop16, Ytm1, Nsa1, Cic1, Rix1, Nop12, Nog2, Arx1 and Lsg1 were used for the affinity purification of ribosome assembly intermediates, each performed in biological triplicates (i.e. pulldowns from independently grown yeast cultures). A wild type *S. cerevisiae* BY4741 strain without any TAP-tagged protein served as control for unspecific binding. In addition, a reciprocal affinity purification with TAP-tagged Cka1 was performed.

For each affinity purification, 12 L YPD medium (2 % (w/v) peptone, 1% (w/v) yeast extract, 2 % (w/v) glucose, 0.002 % (w/v) adenine) were inoculated from 300 ml over-night culture with an OD_600_ of 0.1 and grown at 30 °C and 130 rpm to OD_600_ = 0.8 – 1.0. Cells were harvested by centrifugation at 4300 x g and 4 °C for 12 min. Cell pellets were resuspended in 80 ml cold lysis buffer (LB-P, 50 mM HEPES pH 7.4, 100 mM KCl, 1.5 mM MgCl_2_, 0.1 % (v/v) NP-40, 5 % (v/v) glycerol, pefabloc 1:100, aprotinin/leupeptin 1:1000) and centrifuged at 4000 x g and 4 °C for 5 min. Washed cell pellets were resuspended in 20 ml LB-P and dripped into liquid nitrogen. Frozen droplets of cell suspension were stored at −80 °C until milling in a pre-cooled Retsch® ball mill MM400 at 30 Hz for 2x 60 s. 150 ml ice cold LB-P was added to the frozen cell powder which was thawed on a rolling mixer at 4 °C. Cell debris was separated from the lysate by centrifugation at 30000 x g and 4 °C for 20 min. The lysate was incubated with 1.2 ml equilibrated IgG sepharose beads (GE Healthcare) at 4 °C for 3 h. IgG beads were washed 3x with LB-P and 1x with LB-DTT (50 mM HEPES pH 7.4, 100 mM KCl, 1.5 mM MgCl_2_, 0.1 % (v/v) NP-40, 5 % (v/v) glycerol, 1 mM DTT). IgG beads were loaded onto a 5 mL Polyprep® column using 3x 10 mL LB-DTT. The column was closed and IgG beads were incubated in 4.5 mL LB-DTT with 175 µl TEV protease (produced in-house, 1.5 µg/µL in 10 % glycerol) at 4 °C over-night on a rolling incubator. IgG eluate was incubated with 1 mL equilibrated calmodulin affinity resin (Agilent) in 15 mL LB-CaCl_2_ (50 mM HEPES pH 7.4, 100 mM KCl, 1.5 mM MgCl_2_, 0.02 % (v/v) NP-40, 5 % (v/v) glycerol, 2 mM CaCl_2_) at a final CaCl_2_ concentration of 2 mM on a rolling mixer at 4 °C for 3 h. Calmodulin beads were loaded onto a 5 mL Polyprep® column using 2x 20 mL LB-CaCl_2_ and washed with 1x 10 mL LB-CaCl_2_. The column was closed and calmodulin beads were incubated with 550 µL LB-EGTA (50 mM HEPES pH 7.4, 100 mM KCl, 1.5 mM MgCl_2_, 0.01 % (v/v) NP-40, 5 % (v/v) glycerol, 5 mM EGTA) for 20 min at 4 °C on a rolling incubator. The eluate was collected and the elution was repeated 3x with 450 µL LB-EGTA. Eluates 1 – 4 were concentrated using an Amicon® Ultra 10 K 0.5 mL filter (Merck Millipore) and the buffer was exchanged to a final volume of ca. 100 µl crosslinking buffer (20 mM HEPES pH 8.3, 5 mM MgCl_2_).

Chemical crosslinking and subsequent analysis were carried out essentially as described previously (Leitner et al., 2014). In short, the isotopically labeled crosslinking reagent disuccinimidyl suberate d0/d12 (DSS-H12/D12, Creativemolecules Inc.) was dissolved in N,N-Dimethylformamide (DMF, Sigma) and purified ribosome assembly intermediates were incubated with DSS-H12/D12 at a final concentration of 1.5 mM for 30 min at 30 °C while shaking at 650 rpm in a Thermomixer (Eppendorf). The reaction was subsequently quenched with ammonium bicarbonate at a final concentration of 50 mM for 10 min at 30 °C and 650 rpm. Crosslinked samples were stored at −20 °C over-night.

### Fractionation and enrichment of crosslinked peptides by size exclusion chromatography

Crosslinked samples were dried (Eppendorf, Concentrator plus), resuspended in 100 µl 8M Urea, reduced, alkylated, and digested with trypsin (Promega). Digested peptides were separated from the solution and retained by a solid phase extraction system (SepPak, Waters). Crosslinked peptides were enriched by size exclusion chromatography using an ÄKTAmicro chromatography system (GE Healthcare) equipped with a Superdex^TM^ Peptide 3.2/30 column (column volume = 2.4 ml). Fractions were collected in 100 µl units and analyzed by LC-MS/MS. For each crosslinked sample three fractions were measured in technical duplicates. The elution fractions 1.0-1.1, 1.1-1.2 and 1.2-1.3 ml containing the largest peptides were pooled and the two elution fractions 1.3-1.4 ml and 1.4-1.5 ml were also analyzed by LC-MS/MS. Absorption levels at 215 nm of each fraction were used to normalize peptide amounts prior to LC-MS/MS analysis.

### LC-MS/MS analysis

LC-MS/MS analysis was carried out on an Orbitrap Fusion Tribrid mass spectrometer (Thermo Electron, San Jose, CA). Peptides were separated on an EASY-nLC 1200 system (Thermo Scientific) at a flow rate of 300 nl/min over an 80 min gradient (5 % acetonitrile in 0.1 % formic acid for 4 min, 5 % − 35 % acetonitrile in 0.1% formic acid in 75 min, 35 % − 80 % acetonitrile in 1 min). Full scan mass spectra were acquired in the Orbitrap at a resolution of 120,000, a scan range of 400 − 1500 m/z, and a maximum injection time of 50 ms. Most intense precursor ions (intensity ≥ 5.0e3) with charge states 3 - 8 and monoisotopic peak determination set to ‘peptide’ were selected for MS/MS fragmentation by CID at 35 % collision energy in a data dependent mode. The duration for dynamic exclusion was set to 60 s. MS/MS spectra were analyzed in the Iontrap at a rapid scan rate.

### Label free quantification and statistical analysis of non-crosslinked peptides

Proteins were identified based on their non-crosslinked peptides and quantified in a label-free approach by MaxQuant (version 1.6.5.0.) using standard settings and the match between runs option (Hein et al., 2015). Detailed search parameters were deposited together with the raw files at PRIDE. As fasta database the *Saccharomyces cerevisiae* (strain ATCC 204508 / S288c) proteome with the proteome ID UP000002311 was downloaded from uniProt on 04.03.2019.

For the downstream analysis of the non-crosslinked proteomic data the Perseus platform (version 1.6.5.0.) was used (Tyanova et al., 2016b). In short, MaxQuant results were filtered for hits in the reverse database, for proteins which have been only identified by site and for potential contaminants. After log2(x) transformation of the LFQ values, proteins were considered to be reproducibly identified, if an LFQ value could be determined in all 3 biological replicates in at least one of the technical replicates of at least one purified pre-ribosomal particle. Missing LFQ values were imputed for the total matrix from a normal distribution (width = 0.3 and down shift = 1.8). In order to test for the variance of the determined LFQ values in the 3 biological replicates of each pre-ribosomal particle, an ANOVA significance test with the parameter s0 = 0 in combination with a permutation-based FDR estimation (FDR = 0.05) was applied. For the permutation-based FDR estimation, technical replicates were indicated and randomized together. Data was normalized by Z-scoring (rows). As a negative control for non-specific binding during the purification of pre-ribosomal particles, a wild type yeast strain containing no TAP-tagged protein was used. If a protein was reproducibly identified with an LFQ value in all 3 biological replicates in at least one of the technical replicates of the negative control, this protein was not considered for further analysis. The remaining 272 proteins were used to hierarchically cluster the different biological and technical replicates of purified pre-ribosomal particles (columns) based on their similarity in protein abundance by an L1 distance metric (**Figure S1**). From the clustering of the pre-ribosomal particles the following timeline for ribosome biogenesis was deduced: Noc1-Rsa3-Ssf1-Nop16-Ytm1-Nsa1-Cic1-Rix1-Nop12-Nog2-Arx1-Lsg1.

In addition, a hierarchical cluster analysis of the 272 proteins (rows) by an L1 metric assisted a rough grouping of these proteins based on their abundance in different pre-ribosomal particles into ‘early’ (red), ‘early-intermediate’ (orange), ‘intermediate’ (blue), intermediate-late’ (yellow) and ‘late’ (green) protein clusters (**Figure 1B, 1C, 1D, 2A & 6A, Supplemental Figures 2 – 4, Supplementary Data 1**).

For some specific subsets of the total of 272 quantified proteins (e.g. the DEADbox helicases and the 4 subunits of Ck2) an additional correlation analysis was carried out, where proteins that showed a similar abundance pattern over the different pre-60S particles were identified. For this purpose, Z-score normalized log2 transformed LFQ values were averaged over the 6 replicate measurements for each of the 12 TAP-tagged bait proteins and a standard correlation coefficient (pearson) was calculated for any two pairs of the 272 proteins. Here, proteins with a correlation coefficient ≥ 0.95 to any of the DEADbox helicases were considered to correlate (**Figure 2C**) and proteins with a correlation coefficient >0.85 to three out of the four subunits of the casein kinase Ck2 complex, one of which had to be ≥ 0.9, were considered to correlate (**Figure 6B**).

### Identification of crosslinked peptides

For the crosslink search, a database containing a total of 384 proteins was compiled including all 117 known r-proteins (including both alleles for homologous r-proteins which do not share 100 % sequence identity), all 81 assembly factors which are present in structures of pre-ribosomal particles as well as all 83 known assembly factors which have not been positioned yet (Woolford and Baserga, 2013). Additionally, the 82 proteins identified in this study as candidate novel RBFs were added to the database for the crosslink search. In the control pulldowns, which were performed from a wild type *S. cerevisiae* BY4741 strain without a TAP-tagged bait protein, 24 proteins were identified. 7 of these 24 proteins were r-proteins or known assembly factors and the remaining 17 proteins were added to the database for the crosslink search. Crosslinks to these 24 proteins were subtracted later during the analysis. In 4 select cases, where proteins count not be unambiguously identified because parts of the sequence could be matched to multiple proteins, additional sequences in order to allow unambiguous assignment were added to the database.

In total, the database for the crosslink search comprised 384 proteins.

MS raw files were converted to centroid files and searched using *xQuest* in ion-tag mode. Crosslinks were exported as .tsv files with the filter settings deltaS = 95 and a max. ppm range from −5 to 5, containing all (non-unique) identifications.

### Mono- and intralink filter (mi-filter) for improved false-discovery rates in medium to large scale AP-XL-MS data

Subsequent filtering and analysis of *xQuest* results was done with python / pandas and jupyter notebook. The FDR was calculated as number of decoys / (number of decoys + number of hits). Please note that crosslinked peptides containing at least 1 decoy_peptide were considered as decoys. Three different filter settings were compared in this study. A “standard” filter setting relying on one identified high-confidence crosslink per unique crosslinking site (uxID) (ld-Score ≥ 25, uxID n=1), a more stringent filter setting relying on two independently identified high-confidence crosslinks per uxID (over all AP-MS datasets) (ld-Score ≥ 25, uxID n=2) and the “final” setting, which was also the final setting used for our analysis, which additionally uses a mono- and intralink filter (mi-filter) which requires that for each protein involved in an inter-protein crosslink also at least one mono- or intralink had to be detected in one of the three biological replicates (ld-Score >/= 25, uxID n=2, mi-filter). After filtering, non-specific crosslinks and non-specific proteins identified in the negative control pulldown (without a bait protein) were subtracted from the datasets. Here, 24 unspecific binders were identified based on non-crosslinked peptides and 7 via crosslinks and therefore subtracted from the datasets.

The final dataset, after filtering and subtraction of unspecific crosslinks / proteins identified in the negative control, contained 43205 monolinks (FDR = 0.00138) consisting of 1362 unique monolink sites (FDR = 0.00365), 15802 intra-protein crosslinks (FDR = 0.00327) consisting of 947 unique intra-link sites (FDR=0.002107) and 2844 inter-protein crosslinks (FDR = 0.00385) consisting of 290 unique crosslinking sites (FDR = 0.0136) and 145 proteins.

### Filtering of crosslinked peptides for reciprocal pulldown on Cka1

The *xQuest* results from 1 biological replicate were exported from the results viewer as .tsv files containing all (non-unique) identifications with the filter settings deltaS = 0.95 and a max. ppm range from −5 to 5. Subsequent filtering and analysis were done with python / pandas and jupyter notebook. Only inter-protein crosslinks between proteins, which were identified with at least 1 mono-, intra-peptide- or intra-protein-link with an ld-Score >/= 25, were considered for further analysis.

### Mapping of filtered crosslinks

Crosslink networks were visualized with xiNet (Combe et al., 2015) and mapped manually onto cryo-EM structures of pre-ribosomal particles. For a better overview, several binding regions of RBFs on the cryo-EM structures were highlighted (*see* **Figure S6**). The crosslink networks of the ‘intermediate’ 60S pre-ribosomal particles Nop16, Ytm1 and Cic1 were laid onto the pdb structure 6ELZ, which was purified as state E sequentially by Rix1-TAP and Rpf2-Flag and contains the Ytm1 E80A mutant for impaired removing of the Erb1-Ytm1 complex by Rea1 (Kater et al., 2017). Crosslink networks of Ssf1 and Nsa1 pre-60S particles were laid onto the earlier state C (pdb 6EM1). The crosslink networks of the ‘intermediate to late’ 60S pre-ribosomal particles Nop12, Nog2 and Arx1 were laid onto the pdb structure 3JCT, which was purified by Nog2 (Wu et al., 2016) and the Rix1 crosslink network was laid onto the pdb structure 6YLH purified by Rix1 and Rea1 (Kater et al., 2020). The crosslink network of Lsg1 was laid onto the pdb structure 6RZZ, which was purified by Lsg1 (Kargas et al., 2019). All crosslink networks (excluding proteins containing only mono- or intralinks) can be seen in **Figures 4 and S7.**

### Data and Software Availability

The MS raw files, MaxQuant search parameters as well as the ‘combined’ results folder of MaxQuant, the crosslink database and original *xQuest* result files and final mi-filtered crosslink data have all been deposited to the ProteomeXchange Consortium via the PRIDE partner repository (Perez-Riverol et al., 2019) with the project accession number PXD021831 (Username; Password: QAs78UBZ).

